# Lightweight compositional analysis of metagenomes with FracMinHash and minimum metagenome covers

**DOI:** 10.1101/2022.01.11.475838

**Authors:** Luiz Irber, Phillip T. Brooks, Taylor Reiter, N. Tessa Pierce-Ward, Mahmudur Rahman Hera, David Koslicki, C. Titus Brown

**Affiliations:** Graduate Group in Computer Science, UC Davis; Department of Population Health and Reproduction, UC Davis; Department of Population Health and Reproduction, UC Davis; Graduate Group in Food Science, UC Davis; Department of Population Health and Reproduction, UC Davis; Department of Computer Science and Engineering, Penn State University; Department of Computer Science and Engineering, Penn State University; Department of Biology, Penn State University; Huck Institutes of the Life Sciences, Penn State University

## Abstract

The identification of reference genomes and taxonomic labels from metagenome data underlies many microbiome studies. Here we describe two algorithms for compositional analysis of metagenome sequencing data. We first investigate the FracMinHash sketching technique, a derivative of modulo hash that supports Jaccard containment estimation between sets of different sizes. We implement FracMinHash in the sourmash software, evaluate its accuracy, and demonstrate large-scale containment searches of metagenomes using 700,000 microbial reference genomes. We next frame shotgun metagenome compositional analysis as the problem of finding a minimum collection of reference genomes that “cover” the known k-mers in a metagenome, a minimum set cover problem. We implement a greedy approximate solution using FracMinHash sketches, and evaluate its accuracy for taxonomic assignment using a CAMI community benchmark. Finally, we show that the minimum metagenome cover can be used to guide the selection of reference genomes for read mapping. sourmash is available as open source software under the BSD 3-Clause license at github.com/dib-lab/sourmash/.

## Introduction

Shotgun DNA sequencing of microbial communities is an important technique for studying host-associated and environmental microbiomes [1,2]. By sampling the genomic content of microbial communities, shotgun metagenomics enables the taxonomic and functional characterization of microbiomes [3,4]. However, this characterization relies critically on the methods and databases used to interpret the sequencing data [5,6,7,8].

Metagenome function and taxonomy are typically inferred from available reference genomes and gene catalogs, via direct genomic alignment [9,10], large-scale protein search [11,12,13], or k-mer matches [14,15]. For many of these methods, the substantial increase in the number of available microbial reference genomes (1.1m in GenBank as of November 2021) presents a significant practical obstacle to comprehensive compositional analyses. Most methods choose representative subsets of available genomic information to analyze; for example, bioBakery 3 provides a database containing 99.2k reference genomes [9]. Scaling metagenome analysis approaches to make use of the rapidly increasing size of GenBank is an active endeavor in the field [16,17].

Here, we describe a lightweight and scalable approach to compositional analysis of shotgun metagenome data based on finding the minimum set of reference genomes that accounts for all known k-mers in a metagenome – a “minimum metagenome cover”. We use a mod-hash based sketching approach for k-mers to reduce memory requirements [18], and implement a polynomial-time greedy approximation algorithm for the minimum set cover analysis [19].

Our approach tackles the selection of appropriate reference genomes for downstream analysis and provides a computationally efficient method for taxonomic classification of metagenome data. Our implementation in the 

~~~
sourmash
~~~

 open source software package works with reference databases containing a million or more microbial genomes and supports multiple taxonomies and private databases.

## Results

We first describe FracMinHash, a sketching technique that supports containment and overlap estimation for DNA sequencing datasets using k-mers. We next frame reference-based metagenome content analysis as the problem of finding a *minimum set cover* for a metagenome using a collection of reference genomes. We then evaluate the accuracy of this approach using a taxonomic classification benchmark. Finally, we demonstrate the utility of this approach by using the genomes from the minimum metagenome cover as reference genomes for read mapping.

### FracMinHash sketches support accurate containment operations

We define the *fractional MinHash*, or FracMinHash as follows: for a hash function ***h*** : **Ω** → [***O*, *H***], on an input set of hash values ***W*** ⊆ **Ω** and for any 0 < = *s* < = ***H***,

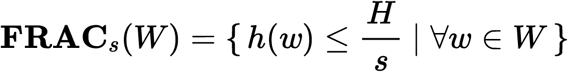

where ***H*** is the largest possible value in the domain of *h*(*x*) and 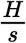 is the *maximum hash value* allowed in the FracMinHash sketch.

The FracMinHash is a mix of MinHash and ModHash [18,20]. It keeps the selection of the smallest elements from MinHash, while using the dynamic size from ModHash to allow containment estimation. However, instead of taking 0 mod *m* elements like **MOD**_*m*_(***W***), a FracMinHash uses the parameter *s* to select a subset of ***W***.

Like ModHash (but not MinHash), FracMinHash supports estimation of the containment index:

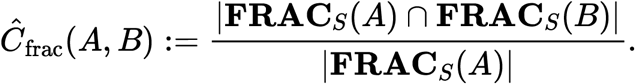

See Methods for details.

Given a uniform hash function *h* and *s* = *m*, the cardinalities of **FRAC**_*s*_(***W***) and **MOD**_*m*_(***W***) converge for large |***W***|. The main difference is the range of possible values in the hash space, since the FracMinHash range is contiguous and the ModHash range is not. This permits a variety of convenient operations on the sketches, including iterative downsampling of FracMinHash sketches as well as conversion to MinHash sketches. Beyond accurate containment operations, FracMinHash can be used to estimate evolutionary distance between pairs of sequences undergoing a mutation model, similar to but more accurately than the MinHash derived method in [20]. See [21] for these and other analytical details.

### A FracMinHash implementation accurately estimates containment between sets of different sizes

We compare the FracMinHash method, implemented in the sourmash software [22] to *Containment MinHash* [23] and Mash Screen (*Containment Score*) [24] for containment queries in data from the 

~~~
podar mock
~~~

 community, a mock bacterial and archaeal community where the reference genomes are largely known [25]; see also Table 1, row 2. This data set has been used in several methods evaluations [24,26,27,28].

**Table 1:**
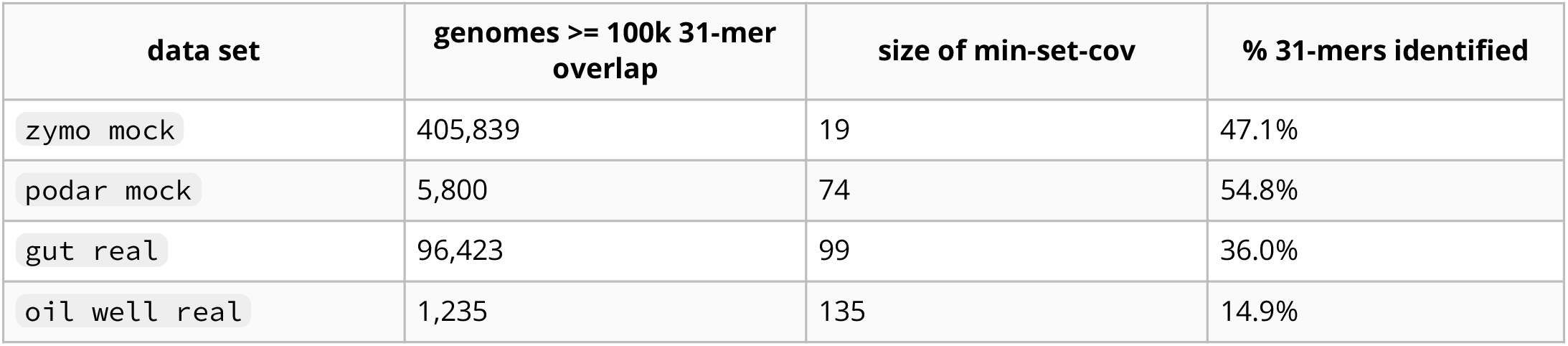
Four metagenomes and the number of genomes in the estimated minimum metagenome cover from GenBank, with scaled=2000 and k=31. Overlap and % 31-mers identified are estimated from FracMinHash sketch size.

| data set | genomes ≥ 100k 31-mer overlap | size of min-set-cov | % 31-mers identified |
| --- | --- | --- | --- |
| zymo mock | 405,839 | 19 | 47.1% |
| podar mock | 5,800 | 74 | 54.8% |
| gut real | 96,423 | 99 | 36.0% |
| oil well real | 1,235 | 135 | 14.9% |

Figure 1 shows containment analysis of genomes in this metagenome, with low-coverage and contaminant genomes (as described in [28] and [24]) removed from the database. All methods are within 1% of the exact containment on average (Figure 1), with 

~~~
CMash
~~~

 consistently underestimating the containment. 

~~~
Mash Screen
~~~

 with ***n*** = **10000** has the smallest difference to ground truth for ***k*** = **{21, 31}**, followed by 

~~~
FracMinHash
~~~

 with 

~~~
scaled=1⊙⊙⊙
~~~

 and 

~~~
Mash Screen
~~~

 with ***n*** = **1000**. The sourmash sketch sizes varied between 431 hashes and 9540 hashes, with a median of 2741 hashes.

**Figure 1:**
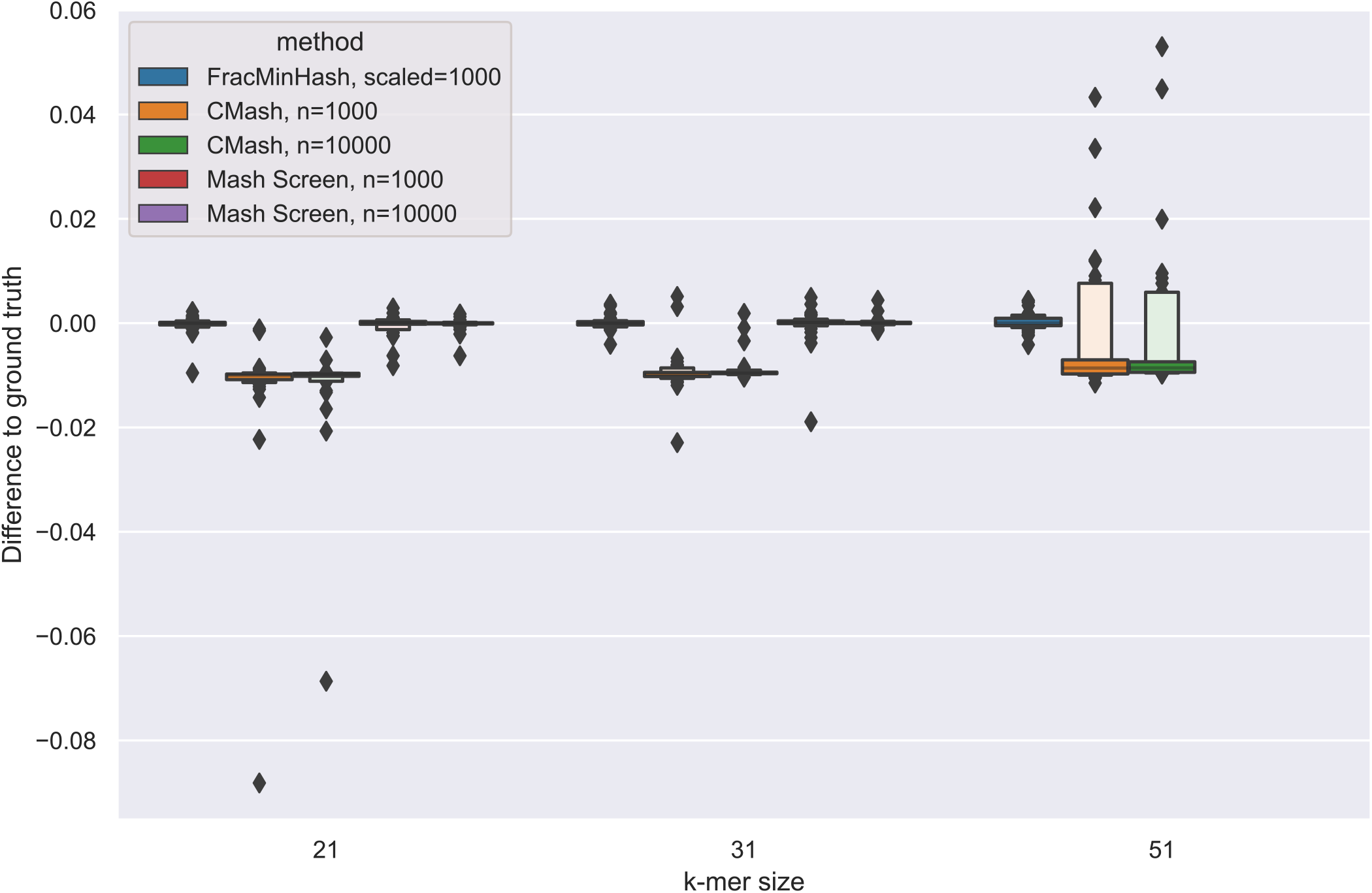
Letter-value plot [29] of the differences from containment estimate to ground truth (exact). Each method is evaluated for ***k*** = **{21, 31, 51}**, except for 

~~~
Mash
~~~

 with ***k*** = **51**, which is unsupported.

### FracMinHash can be used to construct a minimum set cover for metagenomes

We next ask: what is the smallest collection of genomes in a database that contains all of the known k-mers in a metagenome? Formally, for a given metagenome ***M*** and a reference database ***D***, what is the minimum collection of genomes in ***D*** which contain all of the k-mers in the intersection of ***D*** and ***M***? We wish to find the smallest set **{*G_n_*}** of genomes in ***D*** such that, for the k-mer decomposition ***k*()**,

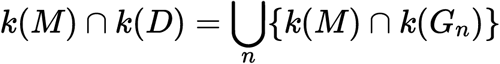

This is a *minimum set covering* problem, for which there is a polynomial-time approximation [19]:

1. Initialize ***C*** ← **∅**
2. While ***k*(*M*)** ∩ ***k*(*D*)** ∖ **⋃**_*G*∈*C*_ (***k*(*M*)** ∩ ***k*(*G*** is: nonempty:
3. ***C*** ← *C* **⋃ {**arg max_*G*∈*D*_ |***k*(*G*)** ∪ (***k*(*M*)** ∩ ***k*(*D*)** ∖ **⋃**_*G*∈*C*_ **(*k*(*M*)** ∪ ***k*(*G*)**))|}
4. return ***C***

This greedy algorithm iteratively chooses reference genomes from ***D*** in order of largest remaining overlap with ***M***, where overlap is in terms of number of k-mers. This results in a progressive classification of the known k-mers in the metagenome to specific genomes.^1^ Note it is classically known that this greedy heuristic results in a logarithmic approximation factor to the optimal set cover solution [19]. This algorithm is implemented as sourmash 

~~~
gather
~~~

.

In Figure 2, we show an example of this progressive classification of 31-mers by matching GenBank genome for 

~~~
podar mock
~~~

. The matching genomes are provided in the order found by the greedy algorithm, i.e. by overlap with remaining k-mers in the metagenome. The high rank (early) matches reflect large and/or mostly-covered genomes with high containment, while later matches reflect genomes that share fewer k-mers with the remaining set of k-mers in the metagenome – smaller genomes, less-covered genomes, and/or genomes with substantial overlap with earlier matches. Where there are overlaps between genomes, shared common k-mers are “claimed” by higher rank matches and only k-mer content specific to the later genome is used to find lower rank matches.

**Figure 2:**
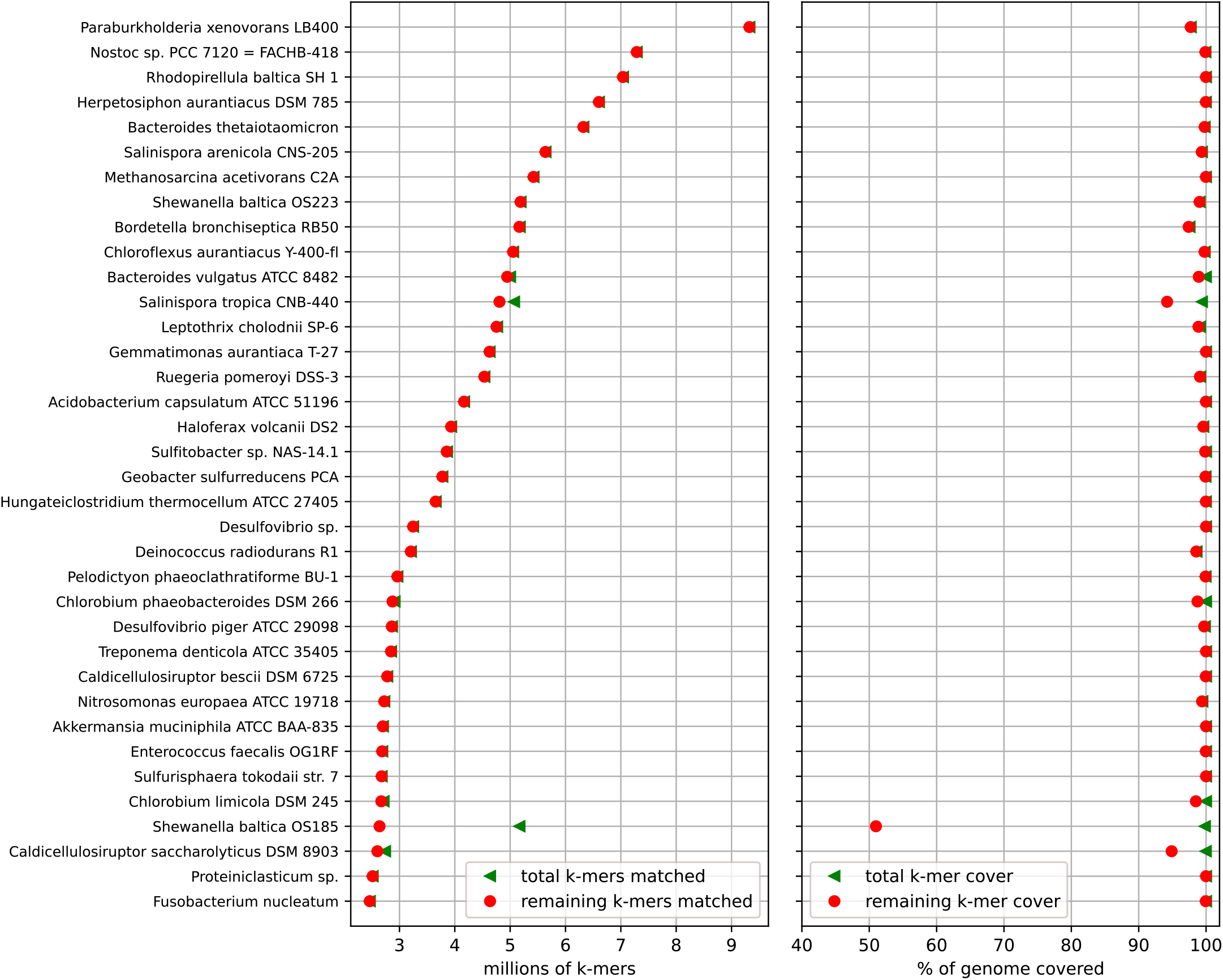
K-mer decomposition of a metagenome into constituent genomes. A rank ordering by remaining containment for the first 36 genomes from the minimum metagenome cover of the 

~~~
podar mock
~~~

 synthetic metagenome [25], calculated using 700,000 genomes from GenBank with scaled=2000, k=31. The Y axis is labeled with the NCBI-designated name of the genome. In the left plot, the X axis represents the estimated number of k-mers shared between each genome and the metagenome. The red circles indicate the number of matching k-mers that were not matched at previous ranks, while the green triangle symbols indicate all matching k-mers. In the right plot, the X axis represents the estimated k-mer coverage of that genome. The red circles indicate the percentage of the genome covered by k-mers remaining at that rank, while the green triangles indicate overlap between the genome and the entire metagenome, including those already assigned at previous ranks.

As one example of metagenome k-mers shared with multiple matches, genomes from two strains of *Shewanella baltica* are present in the mock metagenome. These genomes overlap in k-mer content by approximately 50%, and these shared k-mers are first claimed by *Shewanella baltica* OS223 – compare *S. baltica* OS223, rank 8, with *S. baltica* OS185, rank 33 in Figure 2. Here the difference between the green triangles (all matched k-mers) and red circles (min-set-cov matched k-mers) for *S. baltica* OS185 represents the k-mers claimed by *S. baltica* OS223.

For this mock metagenome, 205m (54.8%) of 375m k-mers were found in GenBank (see Table 1, row 2). The remaining 169m (45.2%) k-mers had no matches, and represent either k-mers introduced by sequencing errors or k-mers from real but unknown community members.

### Minimum metagenome covers can accurately estimate taxonomic composition

We evaluated the accuracy of min-set-cov for metagenome decomposition using benchmarks from the Critical Assessment of Metagenome Interpretation (CAMI), a community-driven initiative for reproducibly benchmarking metagenomic methods [30]. We used the mouse gut metagenome dataset [31], in which a simulated mouse gut metagenome (*MGM*) was derived from 791 bacterial and archaeal genomes, representing 8 phyla, 18 classes, 26 orders, 50 families, 157 genera, and 549 species. Sixty-four samples were generated with *CAMISIM*, with 91.8 genomes present in each sample on average. Each sample is 5 GB in size, and both short-read (Illumina) and long-read (PacBio) simulated sequencing data is available.

Since min-set-cov yields only a collection of genomes, this collection must be converted into a taxonomy with relative abundances for benchmarking with CAMI. We developed the following procedure for generating a taxonomic profile from a given metagenome cover. For each genome match, we note the species designation in the NCBI taxonomy for that genome. Then, we calculate the fraction of the genome remaining in the metagenome after k-mers belonging to higher-rank genomes have been removed (i.e. red circles in Figure 2 (a)). We multiply this fraction by the median abundance of the hashes in the sketch to weight the contribution of the genome’s species designation to the metagenome taxonomy. This procedure produces an estimate of that species’ taxonomic contribution to the metagenome, normalized by the genome size.

In Figures 3 and 4 we show an updated version of Figure 6 from [31] that includes our method, implemented in the 

~~~
sourmash
~~~

 software and benchmarked using OPAL [32]. The minimum metagenome cover was calculated against the Jan 8, 2019 snapshot of RefSeq provided by the CAMI project. Here we compare 10 different methods for taxonomic profiling and their characteristics at each taxonomic rank. While previous methods show reduced completeness – the ratio of taxa correctly identified in the ground truth – below the genus level, 

~~~
sourmash
~~~

 can reach 88.7% completeness at the species level with the highest purity (the ratio of correctly predicted taxa over all predicted taxa) across all methods: 95.9% when filtering predictions below 1% abundance, and 97% for unfiltered results. 

~~~
sourmash
~~~

 also has the second lowest L1-norm error, the highest number of true positives and the lowest number of false positives.

**Figure 3:**
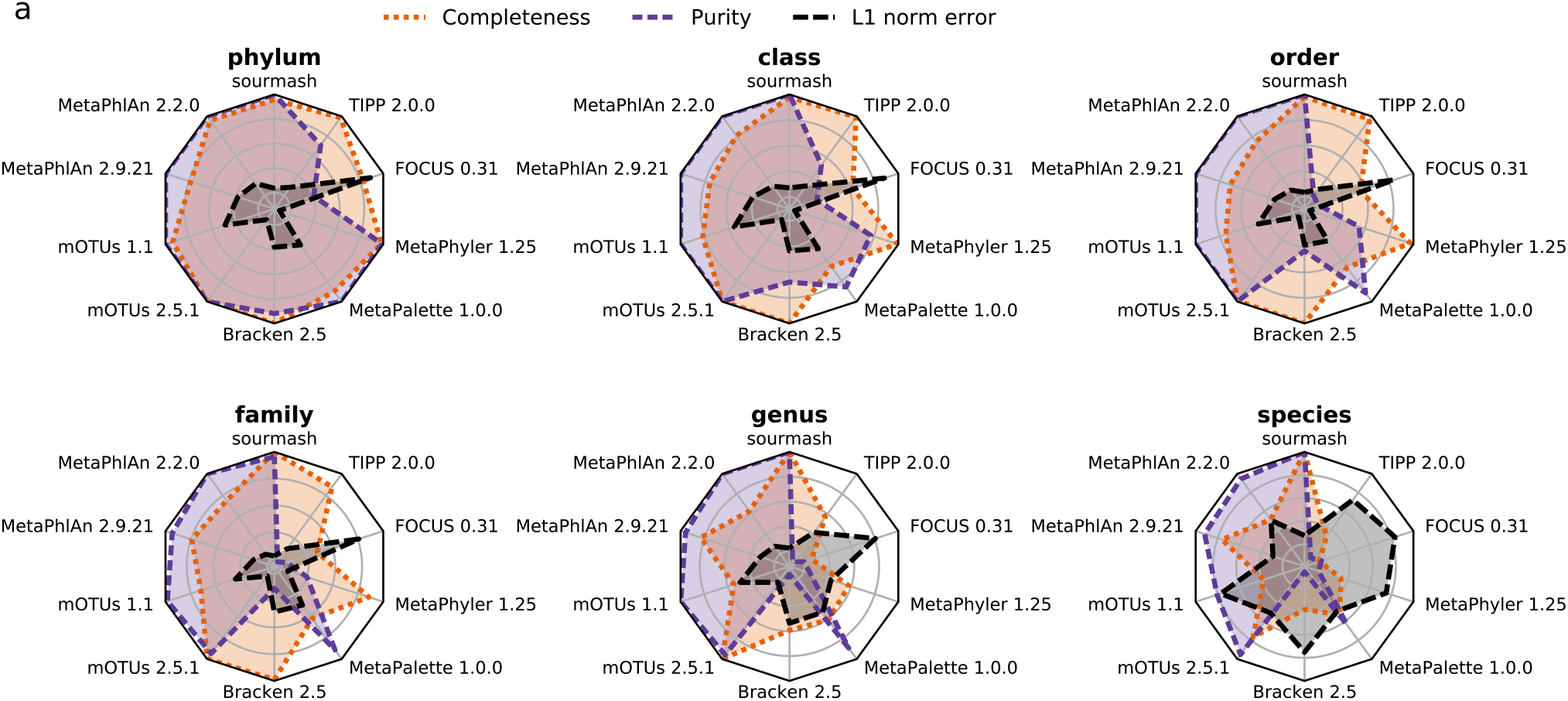
Comparison per taxonomic rank of methods in terms of completeness, purity (1% filtered), and L1 norm.

**Figure 4:**
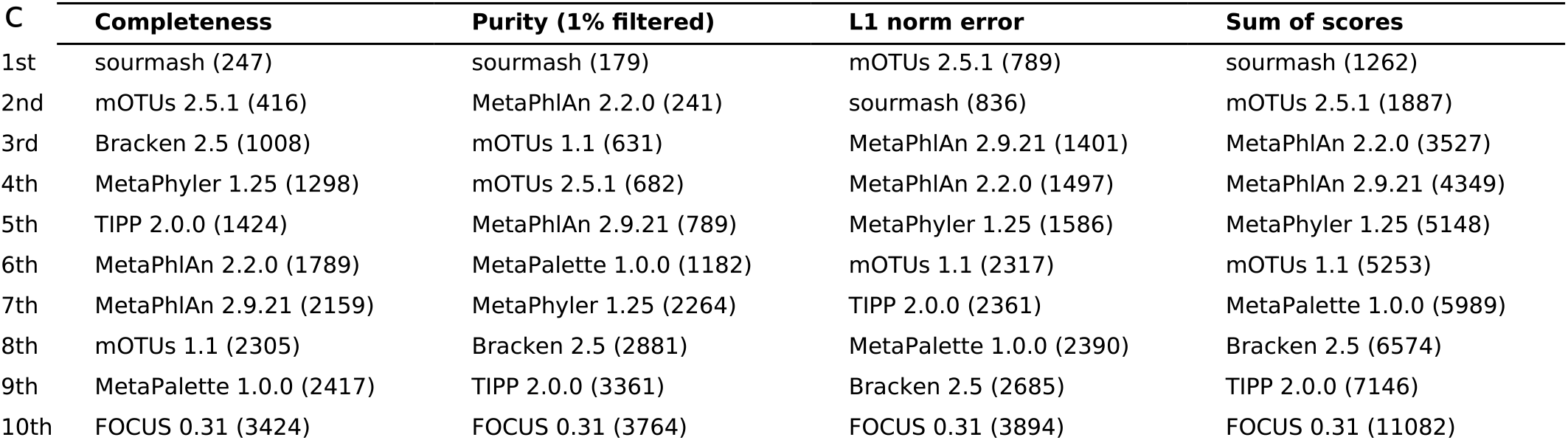
Methods rankings and scores obtained for the different metrics over all samples and taxonomic ranks. For score calculation, all metrics were weighted equally. A scaled value of 2000 and a k-mer size of 31 was used.

### Minimum metagenome covers select small subsets of large databases

In Table 1, we show the minimum metagenome cover for four metagenomes against GenBank - two mock communities [25,33], a human gut microbiome data set from iHMP [3], and an oil well sample [34]. Our implementation provides estimates for both the *total* number of genomes with substantial overlap to a query genome, and the minimum set of genomes that account for k-mers with overlap in the query metagenome. Note that only matches estimated to have more than 100,000 overlapping k-mers are shown (see Methods for details).

We find many genomes with overlaps for each metagenome, due to the redundancy of the reference database. For example, 

~~~
zymo mock
~~~

 contains a *Salmonella* genome, and there are over 200,000 *Salmonella* genomes that match to it in GenBank. Likewise, 

~~~
gut real
~~~

 matches to over 75,000 *E. coli* genomes in GenBank. Since neither 

~~~
podar mock
~~~

 nor 

~~~
oil well real
~~~

 contain genomes from species with substantial representation in GenBank, they yield many fewer total overlapping genomes.

Regardless of the number of genomes in the database with substantial overlap, the estimated *minimum* collection of genomes is always much smaller than the number of genomes with overlaps. In the cases where the k-mers in the metagenome are mostly identified, this is because of database redundancy: e.g. in the case of 

~~~
zymo mock
~~~

, the min-set-cov algorithm chooses precisely one *Salmonella* genome from the 200,000+ available. Conversely, in the case of 

~~~
oil well real
~~~

, much of the sample is not identified, suggesting that the small size of the covering set is because much of the sample is not represented in the database.

### Minimum metagenome covers provide representative genomes for mapping

Mapping metagenome reads to representative genomes is an important step in many microbiome analysis pipelines, but mapping approaches struggle with large, redundant databases [16,17]. One specific use for a minimum metagenome cover could be to select a small set of representative genomes for mapping. We therefore developed a hybrid selection and mapping pipeline that uses the rank-ordered min-set-cov results to map reads to candidate genomes.

We first map all metagenome reads to the first ranked genome in the minimum metagenome cover, and then remove successfully mapped reads from the metagenome. Remaining unmapped reads are then mapped to the second rank genome, and this then continues until all genomes have been used. That is, all reads mapped to the rank-1 genome in Figure 2 are removed from the rank-2 genome mapping, and all reads mapping to rank-1 and rank-2 genomes are removed from the rank-3 genome mapping, and so on. This produces results directly analogous to those presented in Figure 2, but for reads rather than k-mers. This approach is implemented in the automated workflow package 

~~~
genome-grist
~~~

; see Methods for details.

Figure 5 compares k-mer assignment rates and mapping rates for the four evaluation metagenomes in Table 1. Broadly speaking, we see that k-mer-based estimates of metagenome composition agree closely with the number of bases covered by mapped reads: the Y axis has not been re-scaled, so k-mer matches and read mapping coverage correspond well. This suggests that the k-mer-based min-set-cov approach effectively selects reference genomes for metagenome read mapping.

**Figure 5:**
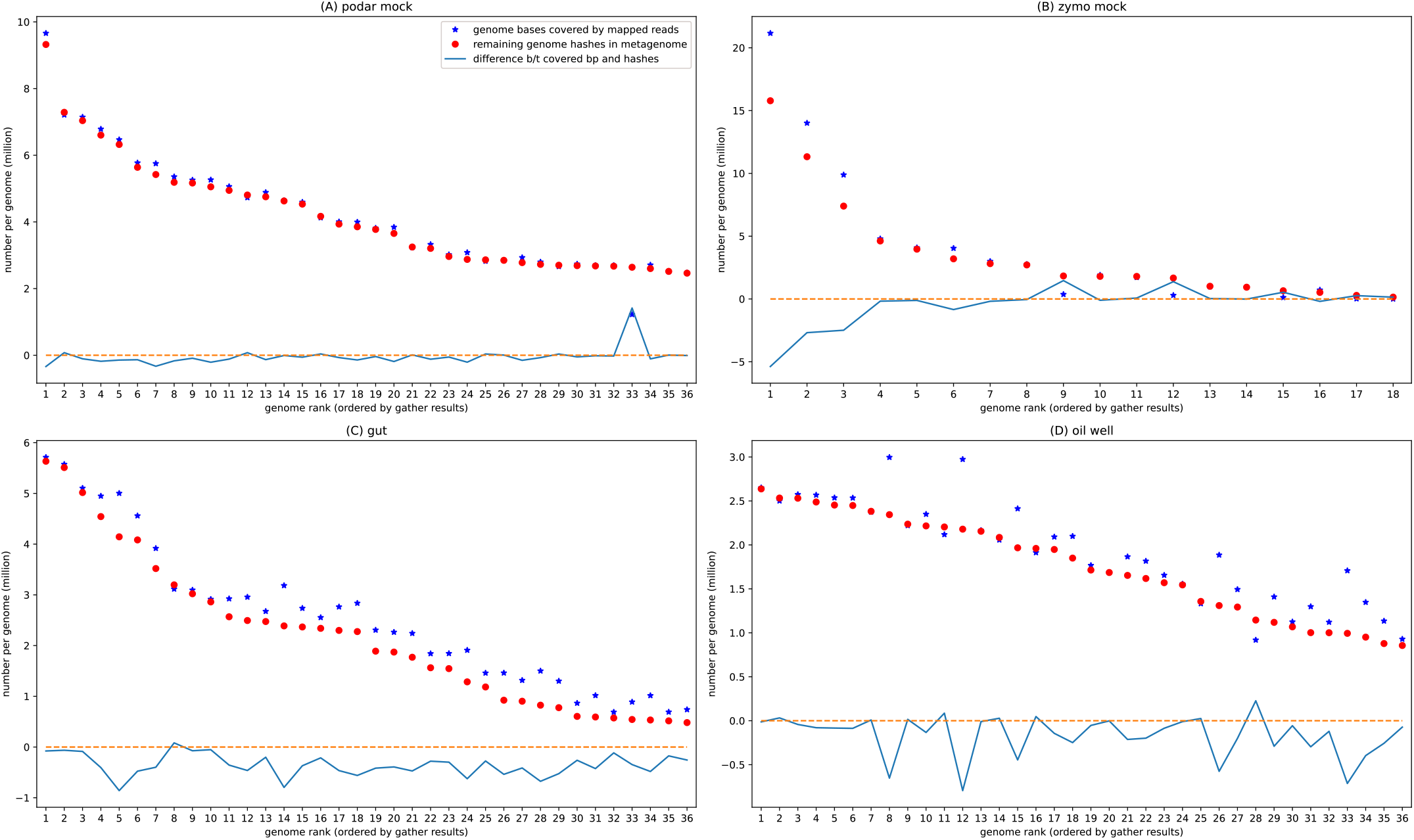
Hash-based k-mer decomposition of a metagenome into constituent genomes compares well to bases covered by read mapping. Plots for each of four metagenomes showing estimated k-mer overlap per genome, along with bases covered by read mapping, for the first 36 genomes in the minimum metagenome cover. The reference genomes are rank ordered along the X axis (as in the Y axis for Figure 2), based on the largest number of hashes from the metagenome specific to that genome; hence the number of hashes classified for each genome (red circles) is monotonically decreasing. The Y axis shows estimated number of k-mers classified to this genome (red circles) or total number of bases in the reference covered by mapped reads (blue stars); the numbers have not been rescaled. Decreases in mapping (peaks in blue lines) occur for genomes which are not exact matches to the genomes of the organisms used to build the mock community; for example, in (A), the peak at rank 33 of 

~~~
podar mock
~~~

 is for *S. baltica OS185*, and represents reads that were preferentially mapped to *S. baltica OS223*, rank 8.

For mock metagenomes (Figure 5 (A) and (B)), there is a close correspondence between mapping and k-mer coverage, while for real metagenomes (Figure 5 (C) and (D)), mapping coverage tends to be higher. This may be because the mock metagenomes are largely constructed from strains with known genomes, so most 31-mers match exactly, while the gut and oil well metagenomes contain a number of strains where only species (and not strain) genomes are present in the database, and so mapping performs better. Further work is needed to evaluate rates of variation across a larger number of metagenomes.

## Discussion

Below, we discuss the use of FracMinHash and minimum metagenome covers to analyze metagenome datasets.

### FracMinHash provides efficient containment queries for large data sets

FracMinHash is a derivative of ModHash that uses the bottom hashing concept from MinHash to support containment operations: all elements in the set to be sketched are hashed, and any hash values below a certain fixed boundary value are kept for the sketch. This fixed boundary value is determined by the desired accuracy for the sketch operations, with clear space/time constraint tradeoffs.

Intuitively, FracMinHash can be viewed as performing density sampling at a rate of 1 ***k***-mer per ***s*** distinct k-mers seen, where *s* is used to define a boundary value 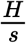 for the bottom sketch. FracMinHash can also be viewed as a type of lossy compression, with a fixed compression ratio of ***s***: for values of ***s*** used here (***s* ≈ 1000**), k-mer sets are reduced in cardinality by 1000-fold.

Unlike MinHash, FracMinHash supports containment estimation between sets of very different sizes, and here we demonstrate that it can be used efficiently and effectively for compositional analysis of shotgun metagenome data sets with k-mers. In particular, FracMinHash is competitive in accuracy with extant MinHash-based techniques for containment analysis, while also supporting Jaccard similarity. In addition, FracMinHash can be used to obtain point estimates of and confidence intervals around mutation rates and evolutionary distances; see [21] for these and other analytical results.

We note that the FracMinHash technique has been used under a number of different names, including Scaled MinHash [35,36], universe minimizers [37], Shasta markers [38], and mincode syncmers [39]. The name FracMinHash was coined by Kristoffer Sahlin in an online discussion on Twitter [40] and chosen by discussants as the least ambiguous option. We use it here accordingly.

FracMinHash offers several conveniences over MinHash. No hash is ever removed from a FracMinHash sketch during construction; thus sketches grow proportionally to the number of distinct k-mers in the sampled data set, but *also* support many operations - including all of the operations used here - without needing to revisit the original data set. This is in contrast to MinHash, which requires auxiliary data structures for many operations - most especially, containment operations [23,24]. Thus FracMinHash sketches serve as compressed indices for the original content for a much broader range of operations than MinHash.

Because FracMinHash sketches collect all hash values below a fixed threshold, they also support streaming analysis of sketches: any operations that used a previously selected value can be cached and updated with newly arriving values. ModHash has similar properties, but this is not the case for MinHash: after ***n*** values are selected any displacement caused by new data can invalidate previous calculations.

FracMinHash also directly supports the addition and subtraction of hash values from a sketch, allowing for limited types of post-processing and filtering without revisiting the original data set. This includes unions and intersections. Although possible for MinHash, in practice this requires oversampling (using a larger ***n***) to account for possibly having fewer than ***n*** values after filtering, e.g. see the approach taken in Finch [41].

When the multiplicity of hashes in the original data is retained, FracMinHash sketches can be filtered on abundance. This allows removing low-abundance values, as implemented in Finch [41]. Filtering values that only appear once was implemented in Mash by using a Bloom filter and only adding values after they were seen once; later versions also implemented an extra counter array to keep track of counts for each value in the MinHash. These operations can be done in FracMinHash without auxiliary data structures.

Another useful operation available on FracMinHash sketches is *downsampling*: the contiguous value range for FracMinHash sketches means that MinHash sketches can be extracted from FracMinHash sketches whenever the size of the requested MinHash is less than the size of the FracMinHash sketch. Likewise, MinHash sketches can be losslessly converted to FracMinHash sketches when the maximum hash value in the MinHash sketch is larger than ***H***/***s***.

Finally, because FracMinHash sketches are simply collections of hashes, existing hash-based k-mer indexing approaches can be applied to sketches to support fast search with both similarity and containment estimators; several index types, including Sequence Bloom Trees [42] and reverse indices, are provided in the sourmash software.

In exchange for these many conveniences, FracMinHash sketches have limited sensitivity for small data sets where the k-mer cardinality of the data set ≈ ***s***, and are only bounded in size by ***H***/***s***, which is typically quite large ≈ **2*e*16**. The limited sensitivity of sketches may affect the sensitivity of gene- and viral genome-sized queries, but at ***s*** = **1000** we see comparable accuracy and sketch size to MinHash for bacterial genome comparisons (Figure 1).

### Minimum set covers can be used for accurate compositional analysis of metagenomes

Many metagenome content analysis approaches use reference genomes to interpret the metagenome content, but most such approaches rely on starting with a list of reduced-redundancy genomes from a much larger database (e.g. bioBakery 3 selects approximately 100,000 genomes [9]), which can reduce sensitivity and precision [17]. Here, we incorporate this reduction into the overall workflow by searching the complete database for a *minimum* set of reference genomes necessary to account for all k-mers shared between the metagenome and the database. We show that this can be resolved efficiently for real-world data sets; implementing a greedy min-set-cov approximation algorithm on top of FracMinHash, we provide an approach that readily scales to 700,000 genomes on current hardware. We show that in practice this procedure reduces the number of genomes under consideration to ≈ **100** for several mock and real metagenomes.

The development of a small list of relevant genomes is particularly useful for large reference databases containing many redundant genomes; for example, in Table 1, we show that for one mock and one real community, we select minimum metagenome covers of 19 and 99 genomes for metagenomes that contain matches to 406k and 96k GenBank genomes total.

The min-set-cov approach for assigning genomes to metagenomes using k-mers differs substantially from extant k-mer and mapping-based approaches for identifying relevant genomes. LCA-based approaches such as Kraken label individual k-mers based on taxonomic lineages in a database, and then use the resulting database of annotated k-mers to assign taxonomy to reads. Mapping- and homology-based approaches such as Diamond use read mapping to genomes or read alignment to gene sequences in order to assign taxonomy and function [43]. These approaches typically focus on assigning *individual* k-mers or reads. In contrast, here we analyze the entire collection of k-mers and assign them *in aggregate* to the *best* genome match, and then repeat until no matches remain.

The resulting minimum metagenome cover can then be used as part of further analyses, including both taxonomic content analysis and read mapping. For taxonomic analyses, we find that this approach is competitive with other current approaches and has several additional conveniences (discussed in detail below). The comparison of hash-based estimation of containment to mapping results in Figure 5 suggests that this approach is an accurate proxy for systematic mapping, as also seen in Metalign [17].

There is one significant drawback to assigning minimum metagenome covers based on k-mers: because k-mers are not a perfect proxy for mapping (e.g. see Figure 5, blue lines), using k-mers to identify the best genome for *mapping* may sometimes lead to inaccurate assignments. Note that long k-mers are generally more stringent and specific than mapping, so e.g. 51-mer overlaps can be used to identify *some* candidate genomes for mapping, but not *all* candidate genomes will necessarily be found using 51-mer overlaps. The extent and impact of this kind of false negative in the min-set-cov approach remains to be evaluated but is likely to only affect strain- and species-level assignments, since nucleotide similarity measures lose sensitivity across more distant taxonomic ranks [44].

Our implementation of the min-set-cov algorithm in sourmash also readily supports using custom reference databases as well as updating minimum metagenome covers with the addition of new reference genomes. When updating metagenome covers with new reference genomes, the first stage of calculating overlaps can be updated with the new genomes (column 2 of Table 1), while the actual calculation of a *minimum* set cover must be redone each time.

Minimum set cover approaches may provide opportunities beyond those discussed here. For example, read- and contig-based analyses, and analysis and presentation of alignments, can be potentially simplified with this approach.

### Minimum metagenome covers support accurate and flexible taxonomic assignment

We can build a taxonomic classifier on top of minimum metagenome covers by reporting the taxonomies of the constituent genomes, weighted by distinct overlap and aggregated at the relevant taxonomic levels. Our CAMI-based taxonomic benchmarking shows that this approach is competitive with several extant approaches for all metrics across all taxonomic levels (Figures 3 and 4). This taxonomic accuracy also suggests that minimum metagenome covers themselves are likely to be accurate, since the taxonomic assignment is built solely on the metagenome cover.

One convenient feature of this approach to taxonomic analysis is that new or changed taxonomies can be readily incorporated by assigning them directly to genome identifiers; the majority of the computational work here is involved in finding the reference genomes, which can have assignments in multiple taxonomic frameworks. For example, sourmash already supports GTDB [45] natively, and will also support the emerging LINS framework [46]. sourmash can also readily incorporate updates to taxonomies, e.g. the frequent updates to the NCBI taxonomy, without requiring expensive reanalysis of the primary metagenome data or even regenerating the minimum metagenome cover.

Interestingly, this framing of taxonomic classification as a minimum set cover problem may also avoid the loss of taxonomic resolution that affects k-mer- and read-based approaches on large databases [47]; this is because we incorporate taxonomy *after* reads and k-mers have been assigned to individual genomes, and choose entire *genomes* based on a greedy best-match-first approach. This minimizes the impact of individual k-mers that may be common to a genus or family, or were mis-assigned as a result of contamination.

Finally, as the underlying min-set-cov implementation supports custom databases, it is straightforward to support taxonomic analysis using *custom* databases and/or custom taxonomic assignments. This is potentially useful for projects that are generating many new genomes and wish to use them for metagenome analysis. sourmash natively supports this functionality.

Our current implementation of taxonomic assignment in sourmash does not provide read-level assignment. However, it is a straightforward (if computationally expensive) exercise to use the read mapping approach developed in this paper to provide read-level taxonomic assignment along with genome abundance estimation.

### The minimum set cover approach is reference dependent

The min-set-cov approach is reference-based, and hence is entirely dependent on the reference database. This may present challenges: for example, in many cases the exact reference strains present in the metagenome will not be present in the database. This manifests in two ways - see Figure 5. First, for real metagenomes, there is a systematic mismatch between the hash content and the mapping content (green line), because mapping software is more permissive in the face of variants than k-mer-based exact matching. Moreover, many of the lower rank genomes in the plot are from the same species but different *strains* as the higher ranked genomes, suggesting that strain-specific portions of the reference are being utilized for matching at lower ranks. In reality, there will usually be a different mixture of strains in the metagenome than is present in the reference database. Methods for updating references from metagenome data sets may provide an opportunity for generating metagenome-specific references [48].

The approach presented here chooses arbitrarily between matches with equivalent numbers of contained k-mers. There are specific genomic circumstances where this approach could usefully be refined with additional criteria. For example, if a phage genome is present in the reference database, and is also present within one or more genomes in the database, it may desirable to select the match with the highest Jaccard *similarity* in order to choose the phage genome. This is algorithmically straightforward to implement when desired.

In light of the strong reference dependence of the min-set-cov approach together with the insensitivity of the FracMinHash technique, it may be useful to explore alternate methods of summarizing the list of overlapping genomes, that is, summarizing *all* the genomes in column 2 of Table 1. For example, a hierarchical approach could be taken to first identify the full list of overlapping genomes using FracMinHash at a low resolution, followed by a higher resolution (but more resource intensive) approach to identify the best matching genomes.

### Opportunities for future improvement of min-set-cov

There are a number of immediate opportunities for future improvement of the min-set-cov approach.

Implementing min-set-cov on top of FracMinHash means our approach may incorrectly choose between very closely related genomes, because the set of subsampled hashes may not discriminate between them. Likewise, the potentially very large size of the sketches may inhibit the application of this approach to very large metagenomes.

These limitations are not intrinsic to min-set-cov, however; any data structure supporting both the *containment* 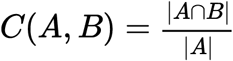 and *remove elements* operations can be used to implement the greedy approximation algorithm. For example, a simple *set* of the *k*-mer composition of the query supports element removal, and calculating containment can be done with regular set operations. Approximate membership query (AMQ) sketches like the Counting Quotient Filter [49] can also be used, with the benefit of reduced storage and memory usage.

In turn, this means that limitations of our current implementation, such as insensitivity to small genomes when *s* is approximately the same as the genome size, may be readily solvable with other sketch types.

There are other opportunities for improving on these initial explorations. The availability of abundance counts for each element in the FracMinHash is not well explored, since the process of *removing elements* from the query does not use them. This may be important for genomes with more repetitive content such as eukaryotic genomes. Both the multiple match as well as the abundance counts issues can benefit from existing solutions taken by other methods, like the *species score* (for disambiguation) and *Expectation-Maximization* (for abundance analysis) approaches from Centrifuge [50].

## Conclusion

The FracMinHash and min-set-cov approaches explored here provide powerful and accurate techniques for analyzing metagenomes, with well defined limitations. We show several immediate applications for both taxonomic and mapping-based analysis of metagenomes. We provide an implementation of these approaches in robust open-source software, together with workflows to enable their practical use on large data sets. The approaches also offer many opportunities for further exploration and improvement with different data structures, alternative approximation algorithms, and additional summarization approaches.

## Methods

### Analytical analysis of FracMinHash

Given two arbitrary sets ***A*** and ***B*** which are subsets of a domain **Ω**, the containment index ***C*(*A*, *B*)** is defined as 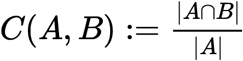. Let ***h*** be a perfect hash function ***h*** : **Ω** → [0, ***H***] for some ***H*** ∈ ℝ. For a *scale factor s* where **0** ≤ ***s*** ≤ **1**, a FracMinHash sketch of a set ***A*** is defined as follows:

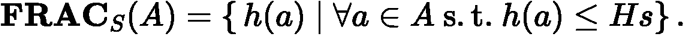

The scale factor ***s*** is a tunable parameter that can modify the size of the sketch. Using this FracMinHash sketch, we define the FracMinHash estimate of the containment index ***Ĉ***_frac_(***A*, *B***) as follows:

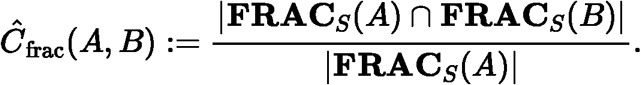

For notational simplicity, we define ***X**_A_* ≔ |**FRAC**_*S*_ (***A***)|. Observe that if one views ***h*** as a uniformly distributed random variable, we have that ***X**_A_* is distributed as a binomial random variable: ***X**_A_* ~ Binom(|***A***|, ***s***). Furthermore, if ***A*** ∩ ***B*** ≠ ∅ where both ***A*** and ***B*** are non-empty sets, then ***X**_A_* and ***X**_B_* are independent when the probability of success is strictly smaller than 1. Using these notations, we compute the expectation of ***Ĉ***_frac_(***A*, *B***).

#### Theorem 1

For 0 < ***s*** < 1, if ***A*** and ***B*** are two distinct sets such that ***A*** ∩ ***B*** is non-empty,

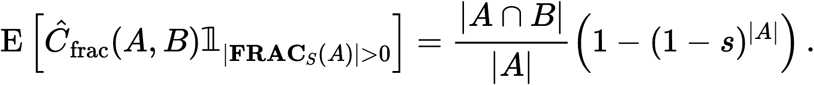

#### proof

Using the notation introduced previously, observe that

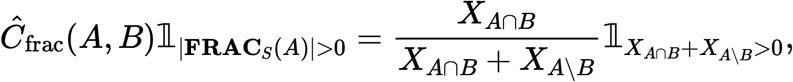

and that the random variables ***X**_**A**_*_∩*B*_ and ***X**_**A**_*_∖***B***_ are independent (which follows directly from the fact that ***A*** ∩ ***B*** is non-empty, and because ***A*** and ***B*** are distinct, ***A*** ∖ ***B*** is also non-empty). We will use the following fact from standard calculus:

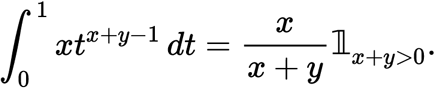

Then using the moment generating function of the binomial distribution, we have

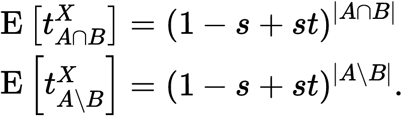

We also know by continuity that

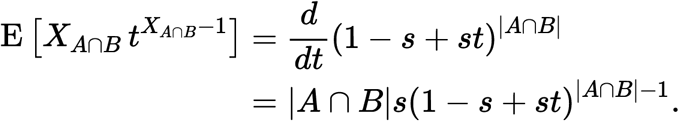

Using these observations, we can then finally calculate that

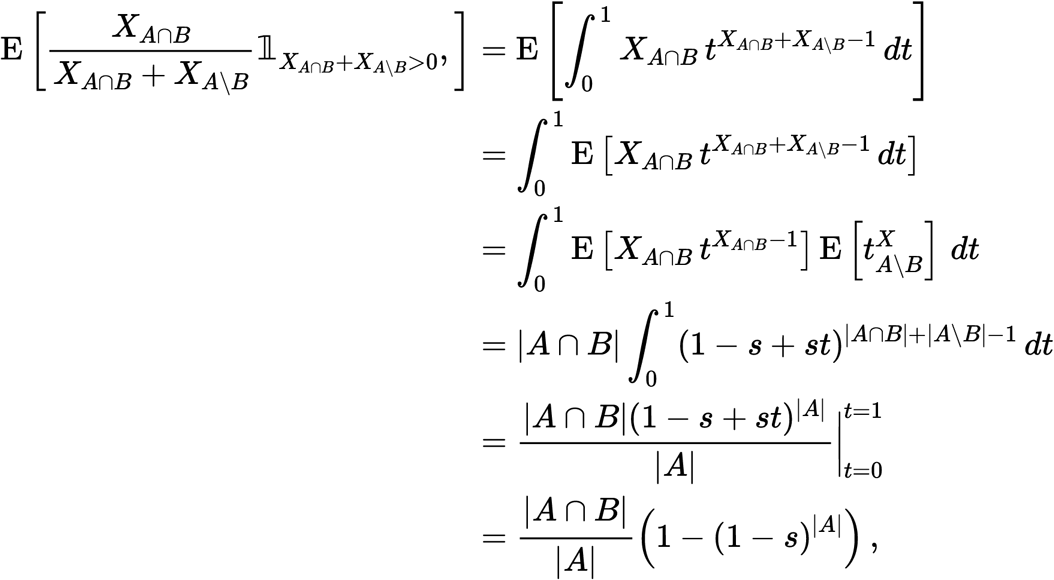

using Fubini’s theorem and independence.

In light of Theorem 1, we note that ***Ĉ***_frac_(***A*, *B***) is *not* an unbiased estimate of ***C***(***A*, *B***). This may explain the observations in [36] that show suboptimal performance for short sequences (e.g. viruses). However, for sufficiently large |***A***| and *s*, the bias factor (1 − (1 − *s*)^|*A*|^) is sufficiently close to 1.

Hence we can define:

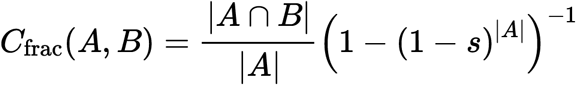

which will have expectation

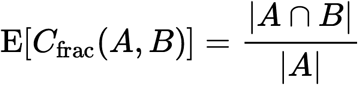

by Theorem 1.

### Implementation of FracMinHash and min-set-cov

We provide implementations of FracMinHash and min-set-cov in the software package 

~~~
sourmash
~~~

, which is implemented in Python and Rust and developed under the BSD license [22]. FracMinHash sketches were created for DNA sequence inputs using the 

~~~
sourmash sketch dna
~~~

 command with the 

~~~
scaled
~~~

 parameter. Minimum metagenome covers were generated using 

~~~
sourmash gather
~~~

 with the sketched metagenome as query against a collection of one or more sketched genomes.

sourmash is available at github.com/sourmash-bio/sourmash. The results in this paper were generated with sourmash v4.2.3.

### Comparison between CMash, mash screen, and Scaled MinHash

Experiments use ***k*** = **{21, 31, 51}** (except for Mash, which only supports ***k*** ≤ **32**). For Mash and CMash they were run with ***n*** = **{1000, 10000}** to evaluate the containment estimates when using larger sketches with sizes comparable to the FracMinHash sketches with ***scaled*** = **1000**. The truth set is calculated using an exact *k*-mer counter implemented with a *HashSet* data structure in the Rust programming language [51]. The sourmash results were generated with 

~~~
sourmash search -- containment
~~~

.

For *Mash Screen* the ratio of hashes matched by total hashes is used instead of the *Containment Score*, since the latter uses a ***k***-mer survival process modeled as a Poisson process first introduced in [52] and later used in the *Mash distance* [20] and *Containment score* [24] formulations.

### GenBank database sketching and searches

Minimum metagenome covers were calculated using a microbial genome subset of GenBank (July 2020, 725,339 genomes) using a scaled factor of 2000 and a k-mer size of 31. Sketches for all genomes and metagenomes were calculated with 

~~~
sourmash sketch dna -p scaled=2⊙⊙⊙,k=31
~~~

. The minimum metagenome covers were calculated using all genomes sharing 50 hashes with the metagenome (that is, an estimated overlap of 100,000 k-mers) with 

~~~
sourmash gather -- threshold-bp 1e5
~~~

. Overlapping sketches were saved with 

~~~
--save-prefetch
~~~

 and matches were saved with 

~~~
--save-matches
~~~

.

The GenBank database used is 24 GB in size and is available for download through the sourmash project [53].

### Taxonomy

The CAMI evaluations were run with the sourmash CAMI pipeline [54], against the Jan 8, 2019 RefSeq snapshot provided by CAMI. This pipeline generated Open-community Profiling Assessment (OPAL) compatible output [30]. This output was then processed with the standard CAMI tools.

### Read mapping and hybrid mapping pipeline

Metagenome reads were mapped to reference genomes using minimap2 v2.17 [55] with short single-end read mapping mode 

~~~
(-x sr)
~~~

.

The hybrid selection and mapping pipeline using the rank-ordered min-set-cov results was implemented in the 

~~~
subtract_gather.py
~~~

 script in the genome-grist package [56].

The complete workflow, from metagenome download to taxonomic analysis and iterative mapping, is implemented in the genome-grist package. genome-grist uses snakemake [57] to define and execute a workflow that combines sourmash sketching, metagenome cover calculation, and taxonomic analysis with metagenome download from the SRA, genome download from GenBank, and read mapping. We used genome-grist v0.7.4 [58] to generate the results in this paper; see 

~~~
conf-paper.yml
~~~

 in the pipeline repository.

genome-grist relies on matplotlib [59], Jupyter Notebook [60], numpy [61], pandas [62], papermill, samtools [63], bedtools [64], fastp [65], khmer [66], screed [67], seqtk [68], and sra-tools [69]. These tools are all installed and managed in snakemake via conda [70] and bioconda [71]. genome-grist itself is developed under the BSD 3-clause open source license, and is available at github.com/dib-lab/genome-grist/.

### Intermediate data products and figure generation

All figures were generated using the Jupyter Notebooks from v0.1 of the github.com/dib-lab/2021-paper-sourmash-gather-pipeline repository [72]. This repository also contains the intermediate data products necessary for figure generation.

### Metagenome data set accessions

The accessions for the metagenome data sets in Table 1 are:

---

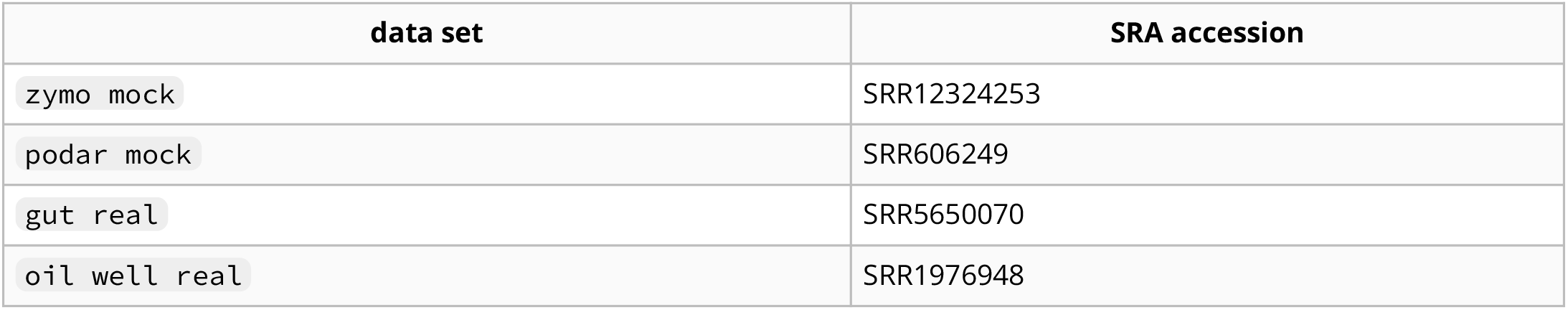

## Footnotes

1. In our current implementation in sourmash, when equivalent matches are available for a given rank, a match is chosen at random. This is an implementation decision that is not intrinsic to the algorithm itself.

